# SIRVs: Spike-In RNA Variants as External Isoform Controls in RNA-Sequencing

**DOI:** 10.1101/080747

**Authors:** Lukas Paul, Petra Kubala, Gudrun Horner, Michael Ante, Igor Holländer, Seitz Alexander, Torsten Reda

**Affiliations:** Lexogen GmbH, Campus Vienna Biocenter 5, 1030 Vienna, Austria

**Author notes:** Correspondence should be addressed to T.R.

**Keywords:** synthetic RNA, *in-vitro* transcripts, IVT, RNA standards, external RNA controls, RNA spike-in mix, RNA sequencing, RNA-Seq, gene expression, sequencing quality control, isoform, transcript variant, poly(A) site

## Abstract

Spike-In RNA variants (SIRVs) enable for the first time the validation of RNA sequencing workflows using external isoform transcript controls. 69 transcripts, derived from seven human model genes, cover the eukaryotic transcriptome complexity of start- and end-site variations, alternative splicing, overlapping genes, and antisense transcription in a condensed format. Reference RNA samples were spiked with SIRV mixes, sequenced, and exemplarily four data evaluation pipelines were challenged to account for biases introduced by the RNA-Seq workflow. The deviations of the respective isoform quantifications from the known inputs allow to determine the comparability of sequencing experiments and to extrapolate to which degree alterations in an RNA-Seq workflow affect gene expression measurements. The SIRVs as external isoform controls are an important gauge for inter-experimental comparability and a modular spike-in contribution to clear the way for diagnostic RNA-Seq applications.

## Introduction

RNA sequencing workflows comprise of RNA purification, library generation, the sequencing itself, and the evaluation of the sequenced fragments (Wang et al. 2009). The first steps impose numerous, whether or not intended, biases towards RNA classes and sequence characteristics, which data processing algorithms try to compensate for afterwards (Li et al. 2010; van Dijk et al. 2014; Zheng et al. 2011). Here, key tasks are the concordant assignment of fragments to the transcript variants, and the subsequent deduction of the corresponding abundance values. As long as the quality of all individual processing steps cannot be unequivocally determined, subsequent comparisons of experimental data, in particular but not exclusively between different data sets, remain ambiguous. The proliferation of different RNA-Seq platforms and protocols has created the need for multi-functional spiked-in controls, which are processed alongside real samples to enable the monitoring and comparing of key performance parameters like the ability to correctly distinguish and quantify transcript variants (Fed. Reg. Doc. 2015-19742).

At present comparisons are carried out only in exemplary inter-laboratory studies on reference RNA samples which investigate different RNA treatments, NGS platforms and data evaluation algorithms (SEQC/MAQC-III Consortium 2014; Li et al. 2014). For these studies reference RNA samples were created from Universal Human Reference RNA (UHRR from Agilent) and Human Brain Reference RNA (HBRR from Ambion, Thermo Fisher), which contain a stable but largely unknown transcript variant diversity. In addition, the reference samples contained also a set of 92 in-vitro transcripts, IVTs, as spike-in controls which were developed by the External RNA Controls Consortium (Baker et al. 2005). These control transcripts, ERCCs (Ambion, Thermo Fisher), allow to asses dynamic range, dose response, lower limit of detection, and fold-change response of RNA sequencing pipelines within the limitation of the mono-exonic, single-isoform RNA sequences (Jiang et al. 2011). Because the ERCCs contain no transcript variants, one of the main challenges of sequencing complex transcriptomes - to identify and distinguish splice variants - could not be evaluated until now.

Here we describe the sequence design and preparation of a comprehensive and novel set of Spike-In Transcript Variants, SIRVs, and show in first examples how mixes thereof can be used to validate isoform-specific RNA sequencing workflows. Further, we demonstrate how to utilize SIRVs for comparing experiments by extrapolating the results from the well-defined ground truth of a small fraction of control reads to the sample reads. Although the focus of the first SIRV module is clearly on the resolution of transcript variants the aim is to build a modular system of IVT controls which retain in the most condensed manner all relevant aspects of the transcriptome complexity to establish a continuous referencing method for RNA sequencing experiments. Then, those controls will provide the base for lasting comparisons of the wealth of RNA-Seq data especially in the face of the rapid development of sequencing technologies.

## Results

### SIRVs Design

The SIRVs were modelled on 7 human genes, whose annotated transcripts were extended by exemplary isoforms to yield between 6 and 18 transcript variants for each gene, and 69 transcripts in total (general design overview in Suppl. Fig. S1). They comprehensively address start- and end-site variations, alternative splicing, overlapping genes, and antisense transcription. Fig. 1A illustrates how the human gene KLK5 served as a blue print for the design of the gene SIRV1 with its 8 transcript variants (gene structures of SIRV2 to SIRV7 are shown in the odd-numbered suppl. Figures from S5 to S15). The SIRV genes model in a condensed and redundant manner all currently known transcription and alternative splicing variations (summary in Fig. 1B). The transcripts range in length from 191 to 2·528 nt (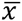, 1·134 nt; 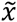, 813 nt), and contain an additional 30 nt long poly(A)-tail. The GC-content varies between 29.5 and 51.2 % (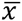, 43.0 %; 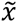, 43.6 %). The exon sequences were created from a pool of database-derived genomes and modified by inverting the sequence to lose identity while maintaining a naturally occurring order in the sequences. The splice junctions conform to 96.9 % to the canonical GT-AG exon-intron junction rule with few exceptions harboring the less frequently occurring variations GC-AG (1.7 %) and AT-AC (0.6 %). Two non-canonical splice sites, CT-AG and CT-AC, account for 0.4 % each. The exon sequences were blasted against the entire NCBI database on the nucleotide and on the protein level and mapped using in silico generated reads. Because no significant matches were found, off-target mapping is de facto absent. Therefore, the artificial SIRV sequences are suitable for non-interfering qualitative and quantitative assessments in known genomic systems, and they are complementary to the ERCC sequences.

**Figure 1.**
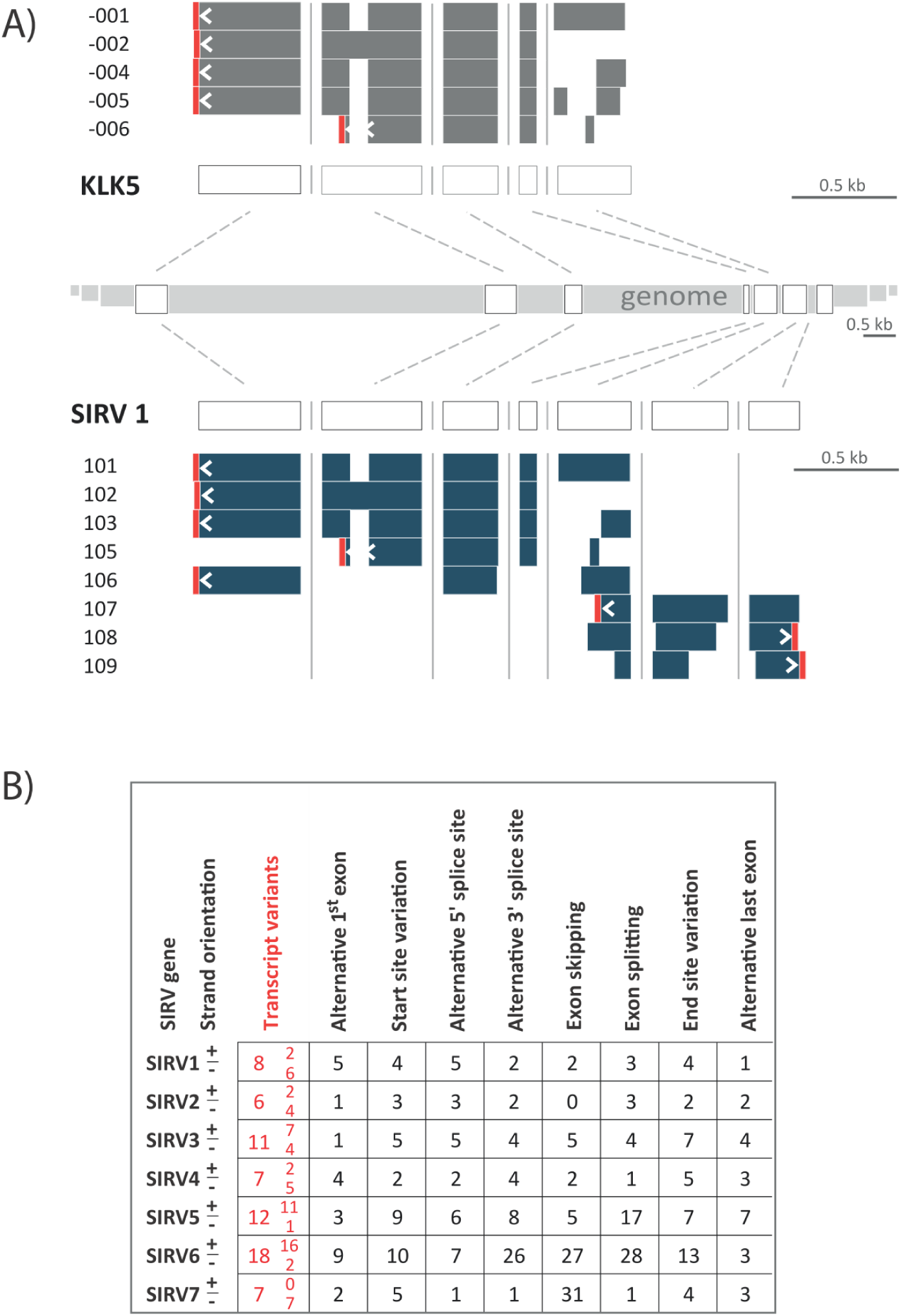
Deriving SIRV1 from model gene KLK5 and classification of implemented transcript isoform features. **A**) Schematic view of the human Kallikrein-related peptidase 5 gene (KLK5) Ensembl annotation (GRCh38.p2) which served as model for SIRV1. The condensed exon-intron structure and the annotated transcript isoforms are shown in the upper grey section. SIRV1 comprises 8 transcript variants, shown in blue. The transcript orientations are indicated by arrows just upstream of the poly(A)-tails highlighted in red. **B)** The occurrences of the different transcription and splicing events are counted for each transcript in reference to a hypothetical master transcript which contains all exon sequences from all transcript variants of a given gene (outlined next to “SIRV 1” in A)). Therefore no intron retention can occur, and exon splitting has been defined to be understood as introduction of an intron sequence.

### Production of SIRV Mixes

The SIRVs were produced by in vitro transcription (IVT) from linearized plasmids, yielding SIRV RNAs of high purity but varying integrity. Since the SIRV transcripts represent isoforms with overlapping sequences, break-down products of SIRV RNAs can potentially affect the detection and quantification of other isoforms. Therefore, a set of tailored purification procedures was applied to obtain full-length RNAs with a minimal amount of side products despite the broad sequence and length variation of the SIRVs (Fig. 2A). By capillary electrophoresis (Bioanalyzer, Agilent) the purified SIRV RNAs were assessed to be at least to 90.31±3.72 % of correct length. Each SIRV transcript entered the final mixes via one of eight PreMixes, allowing for the unique identification of each SIRV by capillary electrophoresis (Fig. 2B). The PreMixes were merged pairwise in equal amounts to yield four SubMixes, which were then combined in defined volumetric ratios to create the final mixes. The experimentally determined cumulative pipetting error for individual SIRVs is smaller than ±8 %, while the applied SubMix scheme limits the maximum error for differential gene expression experiments to only ±4 %. Bioanalyzer traces were used to monitor the relative propagation of the SIRVs in Pre- and SubMixes during the staged mixing (Fig. 2B). The mix E0 maintains equimolar SIRV concentration ratios; E1 contains a maximum concentration difference of 8-fold (close to one order of magnitude); and E2 comprises a concentration difference of up to 128-fold. The inter-mix concentration ratios range from 1/64- to 16-fold (Fig. 2C).

**Figure 2.**
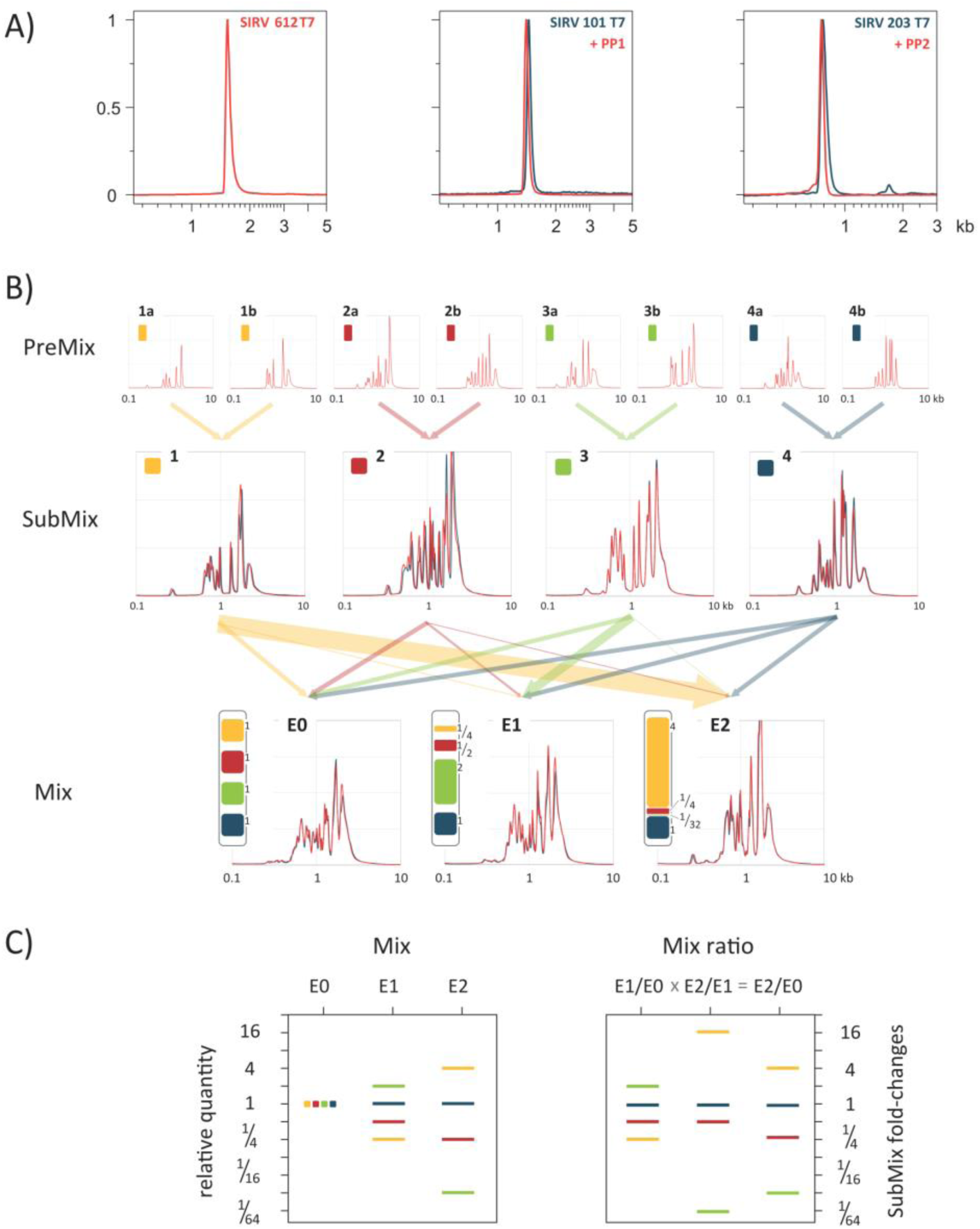
SIRV RNA integrity and mixing scheme. **A**) Examples of high SIRV integrity. T7 transcription of SIRV612 yielded RNA of almost uniform correct length with 4/92/4 % in the pre-/ main-/ and post-peak fractions as analyzed by Bioanalyzer RNA 6000 Pico Chip (Agilent) capillary electrophoresis. The lower integrity of SIRV101 and SIRV203 after T7 transcription (blue traces) was increased using tailored purification protocols, PP1 and PP2 (red traces). Finally, the integrity of all SIRVs was ≥85 wt% with on average 90.3±3.7 wt% in the main peak fraction. **B)** 8 PreMixes received in equimolar entities between 6 and 11 SIRVs which varied in length to be unambiguously traced in Bioanalyzer runs. PreMixes were combined pairwise in equal ratios to yield SubMixes. The resulting 4 SubMixes were combined in defined ratios to obtain the final Mixes E0, E1 and E2. Measured traces are shown in red, and traces computed from the PreMix traces to validate the composition of the SubMixes and final Mixes are shown in blue. A color code for the SubMixes is introduced to follow groupings of SIRVs throughout the subsequent data analysis. **C)** Graphical representation of the relative SIRV concentrations and concentration ratios according their SubMix belonging. The total molarity and weight of the final mixes are evenly balanced at 69.7 fmol/µl and 25.3 ng/µl.

Three different samples with SIRVs, RC-0, RC-1 and RC-2 (for details see methods section), were prepared, NGS libraries generated, and sequenced in paired-end 125 bp mode.

### NGS Data Evaluation

All demultiplexed NGS reads were mapped to the human genome (GRCh38.p2), the ERCC sequences and the SIRVome by using the common splice-aware aligners STAR (Dobin et al. 2012), TopHat2 (Kim et al. 2013), while Bowtie2 (Langmead and Salzberg 2012) was used to map reads to a set of accordingly generated transcript sequences. Bowtie2 aligned reads were further processed by RSEM (Li and Dewey 2011), a combination referred to below as data evaluation pipeline **P1**. Salmon is an autarkic program algorithm (Patro et al. 2015), referred to as **P2**. STAR as well as TopHat2 aligned reads were further processed by Cufflinks2 (Trapnell et al. 2012), and resemble the evaluation pipelines **P3**, and **P4**.

### Estimation of the mRNA Contents

On the basis of the assumption that the RNAs in the samples and the spike-in controls are equally targeted by the library preparation due to similar length distributions (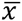_SIRV_ 1·129±15 nt, 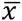_ERCC_ 880±26 nt, and based on the P4 results 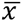_UHRR_ 1·538 nt, and 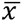_HBRR_ 1·641 nt) the relative mass partition between controls and endogenous RNA can be determined. The propagation of the input amounts to the output read ratio depends on the mRNA content and the relative recovery efficiencies of controls and mRNA after the library preparation. Therefore, one reference value needs to be defined or externally determined, which we chose to be an mRNA content of 3 % in UHRR (Shippy et al. 2006). We further set the efficiency of the endogenous mRNA proceeding through this particular library preparation to 100 %. The SIRVs can then be calculated to progress with a relative efficiency of 87.0±6 %, allowing to determine the mRNA content in the second sample, HBRR, to be 1.72 %. The mRNA content in the mixed samples RC-2 is then assessed to be 2.34 % which corresponds to a minor −9.1 % offset relative to the theoretical outcome based on the RC-0 and RC-1 measurements. In the TruSeq library preparation, the 30 nt long poly(A)-tail of the SIRVs ensures an enrichment similar to the endogenous mRNAs. The ERCC controls show a lower recovery efficiency of 41.6±4.7 % most likely due to their shorter poly(A)tails which vary between 19 and 25 adenines (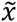, 23 nt). The results are summarized in suppl. Tab. S1.

### Comparison Between Expected and Measured Coverage and Concentration Measures

For the first time the ground truth of complex input sequences is available allowing detailed target-performance comparisons for read alignment, relative abundance calculation, and differential expression determination. Fig. 3 shows for one example, gene SIRV 3 in E0, the expected coverages and counts of the telling junctions together with the coverages obtained by STAR and TopHat2 mapping. NGS workflow-specific read start-site distributions lead to coverage patterns with inherent terminal deficiencies for which the expected coverage has been adjusted accordingly. However, in the measured coverages these systematic start- and end-site biases are accompanied by a variety of biases which introduce severe local deviations from the expected coverage.

**Figure 3.**
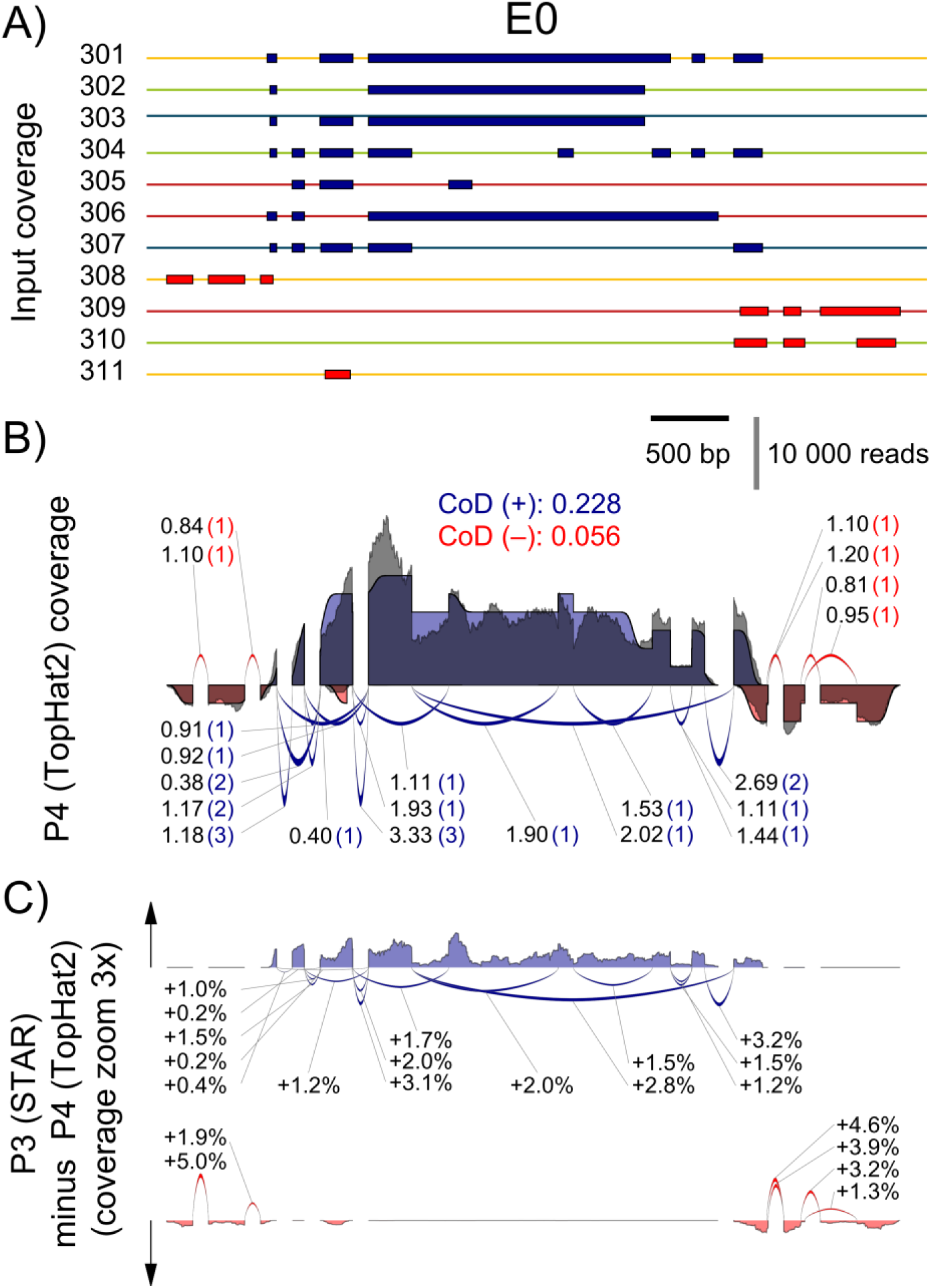
Comparison of the expected and the measured coverages for the SIRV3 locus in mix E0. **A**) Individual transcripts of SIRV3 with transcripts on the plus strand in blue and in red for the ones on the minus strand. Color code indicates the SubMix allocation. **B)** The expected SIRV3 coverage is shown as superposition of individual transcript coverages, in which the terminal sites have been modelled by a transient error function. The measured coverages after read mapping by TopHat2 are shown in grey. The measured coverages and number of splice junction reads were normalized to obtained identical areas under the curves and identical sums of all junctions for the expected and measured data. The measured splice junction reads are shown by the numbers before the brackets, while the expected values are shown inside the brackets. The CoD values are given for the plus and minus strand in the respective colors. **C)** Differences between mapping by STAR aligner and TopHat2. The black arrows indicate the direction of positive differences. Percentages refer to the relative changes of the measured splice junction reads

To obtain a comparative measure, gene-specific coefficients of deviation, CoD, were calculated. CoDs describe the often hidden biases in the sequence data predominantly caused by an inhomogeneous library preparation, but also by the subsequent sequencing and mapping. In the presented example, the stranded TruSeq library preparation with the Illumina HiSeq2500 sequencing and mapping by TopHat2, results in a mean CoD of 0.190±0.193 across all SIRVs. For the sense oriented SIRVs the mean CoD is 0.275±0.223, and for the antisense SIRVs, which cover a region of fewer overlapping transcript annotations, the mean CoD is 0.090±0.076. The coverage target-performance comparison highlights the inherent difficulties in deconvoluting read distributions to correctly identify transcript variants and determine concentrations. The distribution of telling reads, splice junctions, and reads towards the termini are references for the assignment of the remaining reads before calculating relative transcript variant abundances. Not surprisingly, although the majority of junction counts correlates well with the expected values, the high CoD values indicate that numerous telling reads deviate significantly within the context of the individual mixes. They also differ between mixes which affects differential expression measurements (see Fig. 3B, and for the coverage of all seven SIRV genes the even suppl. Fig. from S4 to S16).

The CoD does not allow to distinguish between periodicity and randomness in the biases nor does it forecast how well a data evaluation strategy can cope with the bias contributions. Nevertheless, smaller CoD values are expected to correlate with a simpler and less error-prone data evaluation. The CoD values can be taken as a first, indicative measure to characterize the mapped data, and to compare data sets for similarity up to this point in the workflow. Target-performance comparisons of all SIRV genes in all three mixes can be found in the supplemental information.

The above described four data analysis pipelines, P1 to P4, were used to calculate transcript abundances for 69 SIRV and 196–165 annotated endogenous transcripts. All obtained results are relative concentration measures yielding either <u>F</u>ragments <u>P</u>er <u>K</u>ilobase of exon per <u>M</u>illion fragments mapped (FPKM) (Mortazavi 2008) or <u>T</u>ranscripts <u>P</u>er <u>M</u>illion (TPM) (Li and Dewey 2011). The concentrations were scaled by linear transformation in such way that all SIRVs together reach 69 in E0, 68.5 in E1, and 70.8 in E2, which is the dimensionless value identical to the fmol/µl in the respective SIRV stock solutions. In consequence, the relative quantities of the SIRVs are compatible to the normalized scale which was introduced in Fig. 2C. Boxplots in Fig. 4 show exemplarily the SIRV concentrations and two discrete differentials as measured by P3 (results from P1, P2, and P4, are shown in suppl. Fig. S17, and standard correlation plots of the four pipelines are shown in suppl. Fig. S18). The majority of all SubMix mean concentration values are in concordance with the expected values. The concentration ratios simulating differential expression measurements are less prone to systematic offsets and show by trend narrower distributions. However, these correlations highlight already the issue of obvious and frequent outliers, i.e. transcript variants that are not well resolved.

**Figure 4.**
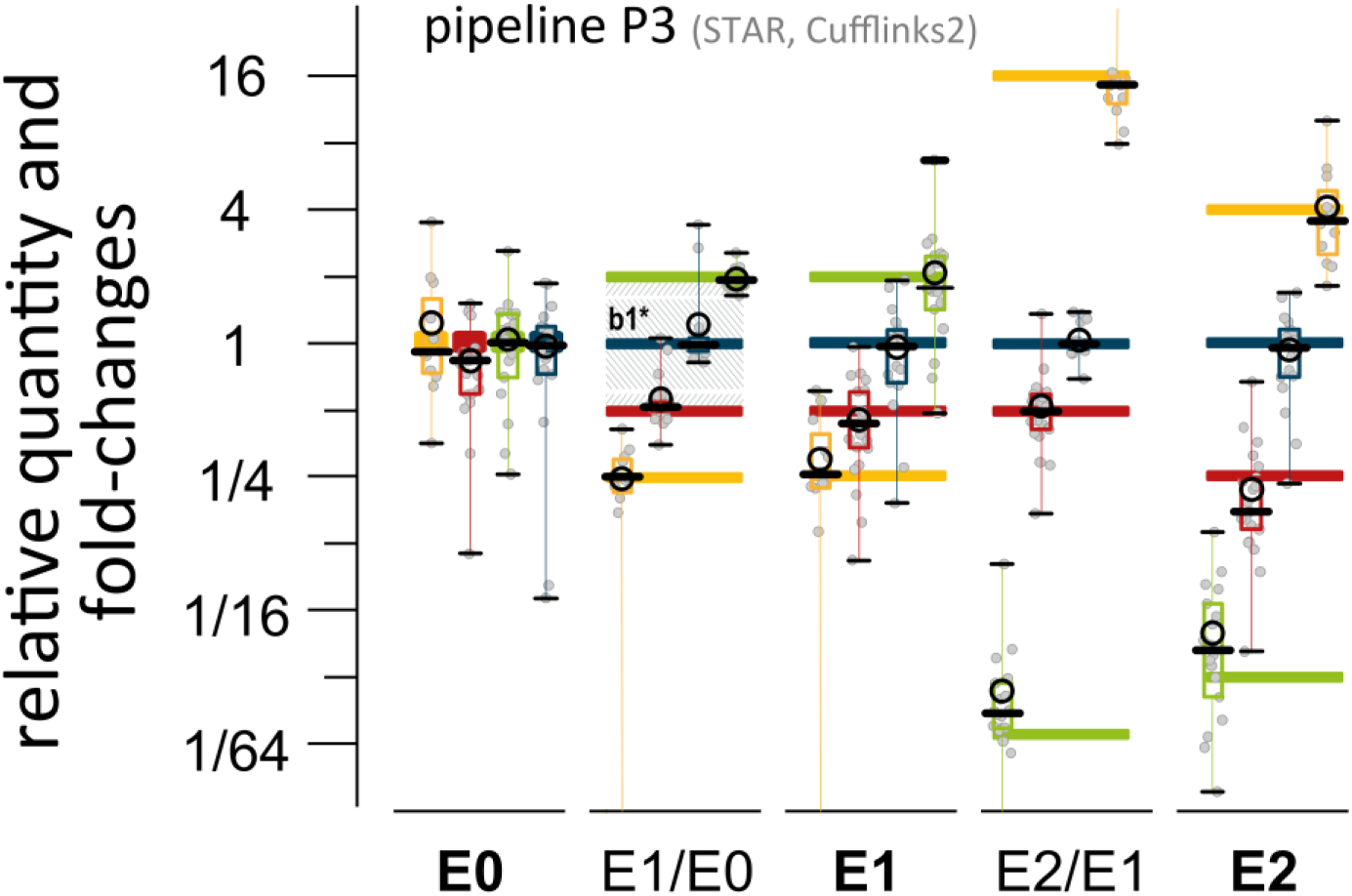
Box plot overview of calculated concentration values for SIRV mixes and mix ratios. Reads from SIRV mixes E0, E1, and E2 were processed by four different data evaluation algorithms for transcript quantification. Results derived by pipeline P3 are exemplarily shown in reference to the known inputs (bars in colors corresponding to the SIRV SubMixes of Fig. 2). The black circles mark the mean and the bold dashes the medians of the data points which are shown in grey, the boxes span the 25 to 75 % region of the data points, and the black whiskers with connecting lines reach up to the min and max values and indicate also outliers outside the scale of the graph. The comparison E1 vs. E0 from sample RC-1 and RC-0 (2^nd^ row) was used for building the histogram of Fig. 5B. The shaded area resembles the 1^st^ bin (0≤ǀLFCǀ≤1), while the inner section (b1*) comprises the fraction 0≤ǀLFCǀ<0.5 where the majority of data points originates from SubMix4 (ratio 1:1) and some “outliers” from SubMix2 which were actually spiked in at the ratio 2:1.

Heat maps allow to inspect each SIRV in the context of competing transcripts. Tab. 1B shows the abundancies as log_2_-fold changes, LFC, relative to the expected values for the example of SIRV 6. LFC values consider relative increases and decreases across all concentration ranges likewise. An LFC window of ±0.11 presents the confidence interval as a result from the currently achievable accuracy in producing the SIRV mixtures. We examine the LFC values uncoupled from the adjusted p-values which provide a statistical measure of the significance level for apparent differential expression. By these means we prevent pipelines to appear more coherent only due to higher noise levels. The *accuracy* has been calculated for each SIRV as the mean of the LFCs in all three mixes. The *precision* is a measure of how consistent SIRVs are quantified, and has been calculated as the LFC standard deviation for each SIRV and, further, for entire pipelines by using all SIRVs. Within the presented example SIRVs can obtain in all four pipelines high precisions of up to 0.18±0.07, and high accuracies of up to −0.01±0.10 (SIRV604). In contrast, SIRV612 of similar length has an average precision of 5.69±3.68 and an accuracy of −3.86±2.70 although it was detected by P2 with a high precision of 0.12 at an accuracy of 0.18. This quantification of differences between pipelines now directly permits to assess the effect of changes in workflows and individual experiments.

### Comparison between Different Workflows

When ranking the four pipelines by said precision P3 gives the lowest value with 1.29, followed by P4 with 1.64, P1 with 1.98, and P2 with 2.14. However, a sample-specific global target-performance comparison is not the strongest criteria for evaluating performance differences at transcript resolution. For this purpose the apparent differential expression shows if similar errors are caused by the same or different transcripts. The highest *concordance* was found between P3 and P1 (labelled P3/P1) calculated as standard deviation of all SIRV LFCs with 1.01, followed by P4/P3 (1.04), P4/P1 (2.55), P2/P1 (2.88), and P3/P2 (3.09). The lowest concordance has P4/P2 with a value of 4.12. One section of the resulting heat map for both extremes is shown in Tab. 1C.

**Table 1.**
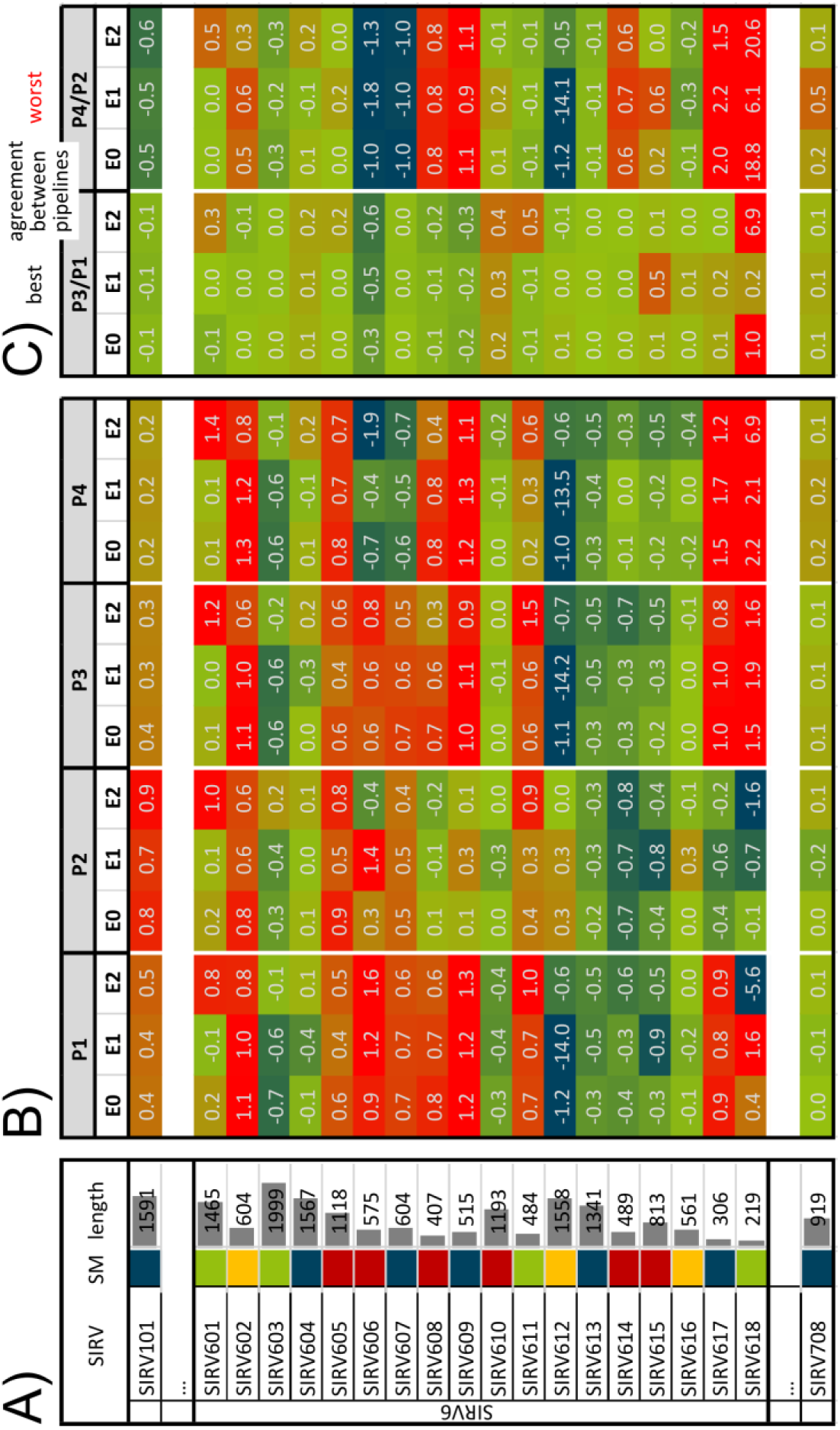
Variance heat map of SIRV concentration measures detailing SIRV6 by the 4 different pipelines P1 to P4. **A)** Given are SIRV transcript identifier, SubMix (SM) color code, and transcript lengths (in nucleotides). **B)** Heat maps showing the log-fold changes (LFC) for the comparison of the normalized expected, and measured SIRV concentrations as a result from the four data evaluation pipelines, P1 to P4, evaluating E0, E1 and E2. Fields with −1 ≤ LFC ≤ 1 are filled with gradient green, all fields with LFC < −1 with straight blue, and all fields with LFC > 1 with straight red. **C)** From all 6 possible pairwise comparisons, P2/P1, P3/P1, … to P4/P3, of the normalized measured SIRV concentrations the one with the smallest (P3/P1) and largest (P4/P2) median of all LFC are shown. The complete heat map is shown in the suppl. Tab. S2.

The same pipeline comparisons were made with either all endogenous transcripts, or the ones in the higher SIRV expression range and compared to the pipeline concordances calculated with the SIRV controls (Fig. 5). We define the first bin with ǀLFCǀ < 1 as the bin of no apparently differentially expressed transcripts independent of the corresponding p-values. The other bins contain transcripts for which the calculated concentrations differ by more than 2-fold, hence convene apparently differentially expressed transcripts (DE). We find that SIRV controls and endogenous transcripts are evaluated coherently. While theoretically no differences are expected, pipelines P3 and P1 differ in their quantification of the same data by 4.4 % for the SIRVs, and by 6.9 % for endogenous RNAs in the SIRV concentration range (Fig. 5A). Pipelines P2 and P4 differ to a larger extent when evaluating the same samples (Fig. 5B, C). The same correlation is seen in comparisons between different samples (shown here for RC-0 vs. RC-1). The expected DE between reference RNAs UHRR and HBRR of approximately 50 % is evaluated differently by P3 vs. P1 (DE of 2.4 %), and this is mirrored in the evaluation of the SIRV mixes spiked into these reference RNAs (DE of 4.9 %). P4 vs. P2 performed worse with a pipeline difference of 3.9 % and 10.3 % for endogenous and SIRV transcripts. The controls show a highly similar increase when comparing P3/P1 with P4/P2 (2.1-fold from 4.9 % to 10.3 %) to the endogenous RNAs (1.6-fold from 2.4 % to 3.9 %). Here, the average measurement using the same pipeline provide a pseudo-ground truth, which is compared to the measurement using different pipelines. It allows to quantify the additional error caused solely by changes in the pipeline. A change from pipeline P3 to P1 would increase the number of DE-detected transcripts to a smaller extent than a change from P2 to P4, and this is true for both, the SIRV controls and the endogenous transcripts.

**Figure 5.**
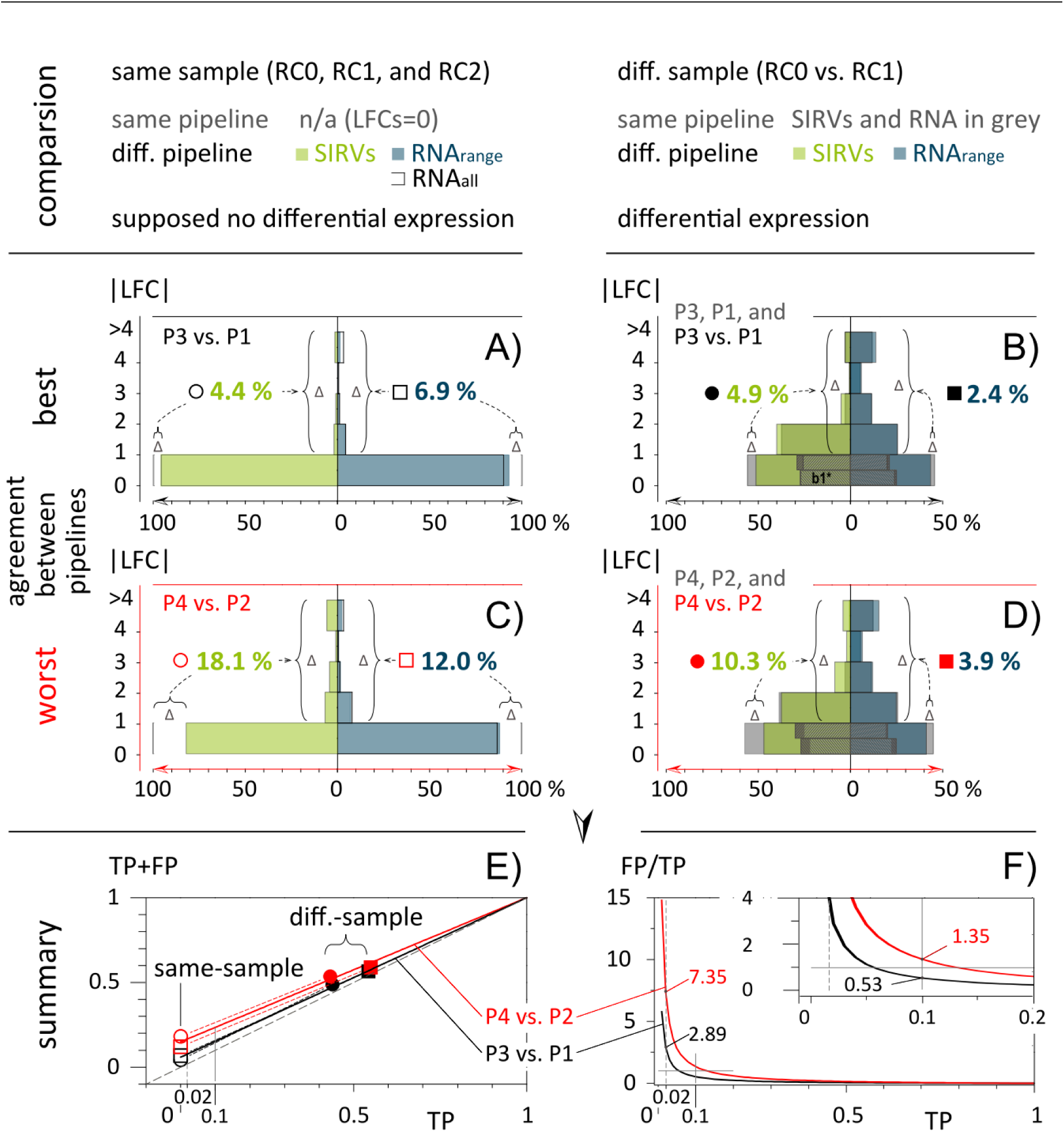
SIRV and endogenous mRNA expression analyses are affected similarly when changing data evaluation pipelines. Binned apparent LFC-moduli are shown in histograms when comparing results obtained by different pipelines evaluating identical NGS data sets from **A, C)** the same samples, or **B, D)** different samples. Exemplarily the comparison with the best, P3 vs. P1 (black symbols), and the worst agreement, P4 vs. P2 (red symbols), are shown. The charts compare side-by-side the results from the SIRV controls (green), all endogenous transcripts (black outline), and endogenous transcripts within the medium-to-high concentration range of 10 to 2250 TPM or FPKM (blue). The average of all intra-pipeline results are shown in grey. The differences, Δ, between analyses with the same and with different pipelines show the losses in the bins 0 < ǀLFCǀ < 1, and the corresponding gain in the other bins ǀLFCǀ > 1. The increase of the FP-rate purely caused by the pipeline changes vs. the TP range is given in the summary graphs **E**, and **F)**. The 1^st^ bins, 0 < ǀLFCǀ < 1, are divided into two 0 < ǀLFCǀ < 0.5, and 0.5 < ǀLFCǀ < 1, which facilitates the easier tracking of the results from the SubMixes4 as shown in Fig. 4, were in both figures the 1^st^ fraction is signed with 1b*.

## Discussion

We report a significant contribution to the external RNA controls toolbox in the form of Spike-In RNA Variants (SIRVs) that address transcription variation and alternative splicing. The highly defined mixes of 69 transcripts aligning to 7 genes enable accurate target-performance evaluations of gene expression analysis methods, which is not possible e.g., by analyzing larger but only loosely defined pools of in vitro transcripts (Lahens et al. 2014), or limited sets addressing special problems like fusion genes (Tembe et al. 2014). The results presented here demonstrate that the condensed complexity of the SIRV controls mimic very closely main properties of endogenous mRNAs when analyzing high-throughput RNA sequencing workflows for their capabilities to determine transcript variants and their relative quantities. The responses of the SIRV controls to changes in the workflows allow to extrapolate measures for the accuracy, precision, and foremost inter-experiment comparability (concordance) to the large pool of endogenous RNAs. With the help of the external controls we could confirm that biases in sequence coverage could mostly be balanced by the exemplary chosen data evaluation algorithms, and for the majority of transcripts consistent relative abundances were obtained. However, in our most concordant same sample pipeline comparison, P3/P1, already 4.4 % of the controls and 6.9 % of the endogenous RNAs in the medium to high concentration range appeared to be differentially expressed although none of those were true positives (TP), because the *reductio ad absurdum* same sample comparison was using the very same sets of NGS reads. In consequence, more than 700 potential endogenous mRNA leads would be non-validatable <u>f</u>alse <u>p</u>ositives (FP) only because of a change in data evaluation pipelines. With a difference between P4 and P2 of 12 % at the level of endogenous RNAs, an extra 1200 FPs would obscure the evaluation.

The inter-sample comparison between UHRR and HBRR presents a rather extreme case where around half of all transcripts were calculated to differ by more than 2-fold. However, the relative FP/TP weight of both comparisons remains similar as P4/P2 shows again clearly higher FPrates in both, controls and endogenous RNAs. The linear interpolation (Fig. 5E) allows to estimate the effect of the experimental changes to relative small number of differences as they would be seen in typical RNA-Seq experiments. Exemplarily highlighted are the intersections at 0.02 and 0.1 in Fig. 5F. TP call rates and FP/TP ratios are the important figures to estimate the likelihood and inversely related effort to validate differentially expressed transcripts. For the P3/P1 comparison at 2 % TP we approximate an FP/TP ratio of 2.89. The likelihood to actually validate DE transcripts under such circumstances would reach 0.35. For the P4/P2 comparison the TP/FP ratio estimates to mere 0.14. The related risks and often substantial costs for deducing potential biomarkers would be 2.5 times higher. Although we demonstrated in the presented example the influence of changes in data evaluation pipeline within an integrated RNA-Seq workflow the same principles apply to the front end the RNA treatment, NGS library preparation, and sequencing. Here, unintended variations in particular sample preparations can only be detected by these external spike-in controls which operate as markers and reference spots. By these means SIRV controls increase the comparability within and between sequencing experiments at the transcript isoform level. SIRVs are designed for the co-development, improvement, and monitoring of library preparations, sequencing, and data processing.

*Outlook:* The presented module and the 92 ERCCs which probe the wide dynamic range using monocistronic transcripts can be stepwise extended in the future by a) ca. 15 poly(A)-tail variations of monocistronic sequences with defined lengths of 20, and 30 adenosines, as well as polyadenylated and purified RNAs with poly(A)-tail lengths close to 50, 100 and above 200 nucleotides, b) around 12 above-average long RNAs with 3, 5, 7, and 10 kb, c) approx. 3 RNAs each with a low, medium, and high GC-content, d) a set of 13 mRNA relevant chemical modifications, most prominent the cap-structure (Bokar and Rottman 1998), e) variants which cover aspects of SNPs, alleles, inversions, and fusion genes, which needs to be accounted for in the corresponding sequence annotation files, and f) a random set of small and micro RNAs that comprise all sequence variations at their termini to account for ligation biases. The external control modules need to be fine-tuned with respect to the dynamic concentration and concentration ratio range, and the reference annotations must provide a mixture of correctly, over-and under-annotated genes to better emulate the challenging conditions for bioinformatical data evaluation. Both optimizations will also further increase the congruence of the presented SIRV module with the endogenous RNAs. Eventually, all modules together would develop into a comprehensive set of approximately 250 long RNA controls plus a microRNA module which will become sufficient for calibration and normalization purposes. This new estimate reduces the scope of designing better spike-in controls compared to the maybe thousands of IVTs which have been previously anticipated (Kratz and Carninci 2014).

## Methods

### SIRV Design and in Silico Analysis

The gene structures of seven human model genes (KLK5, LDHD, LGALS17A, DAPK3, HAUS5, USF2, and TESK2) were used as scaffold for the design of SIRV1 to SIRV7. The ENCODE-annotated transcripts as well as added variants were edited to represent in a redundant and comprehensive manner known transcription and alternative splicing variations (complete gene structures are shown in the odd-numbered suppl. Fig. from S3 to S15). All exon sequences descend from a pool of database-derived genomes (gene fragments from viruses and bacteriophage capsid proteins and glycoproteins selected based on a GC-content in a range of 30-50%) which were modified by inverting the sequence to lose identity while maintaining a kind of naturally occurring order in the sequences. Intron sequences that do not align with exons of another isoform were drawn from random sequences whereby the GC content was balanced to comply with the adjacent exonic sequences (Random DNA Sequence Generator, http://www.faculty.ucr.edu/~mmaduro/random.htm).

To verify the sequence exclusivity, first, the exon sequences were blasted against the entire NCBI database including ERCCs on the nucleotide and protein level (program BLASTN 2.2.29, https://blast.ncbi.nlm.nih.gov/). The default BLAST parameters used were: word size 28, expected threshold 10, Max matches in query rang 20, Match/Mismatch Score 1, −2, Gap Costs Linear, Filter: Low complexity regions, Mask for lookup table only. Search of the nucleotide collection resulted for all SIRVs in “no significant similarity found”. Second, in silico generated reads (Griebel et al. 2012) (FLUX-Generator, http://sammeth.net/confluence/display/SIM/Home, command line: /flux-simulator-p~/…/Flux/sirv1.par -t simulator -x -l -s) were mapped to nine individually selected genomes using TopHat2 (Kim et al. 2013) and Bowtie2 (Langmead and Salzberg2012). Because Bowtie2-aligned reads can be as short as 20 nt, the resulting limited number of mapping reads would be deemed to be not significant in standard NGS work-flows. In contrast, when mapping reads with the standard setting of the splicing-aware TopHat2 program, not a single read aligned to any of the tested genomes. Given that both searches found no significant matches, off-target mapping of the SIRVs can essentially be ruled out. Therefore the artificial SIRV sequences are suitable for noninterfering qualitative and quantitative assessments in the context of known genomic systems and complementary to the ERCC sequences (National Institute of Standards and Technology (NIST) certified DNA plasmids, Standard Reference Material^®^ 2374). The SIRV sequences can be found at https://www.lexogen.com/sirvs/#sirvsdownloads, in the Source Data files, and they are also deposited at the NCBI Nucleotide Database (PRJNA312485).

### SIRV Constructs and the Production of in-vitro Transcripts

Synthetic gene constructs were produced (Bio Basic Inc, Markham, ON, Canada) that comprised 5’ to 3’ a unique restriction site, a T7 RNA polymerase promoter whose 3’ G is the first nucleotide of the actual SIRV sequence, which is seamlessly followed by a (A_30_)-tail that is fused with an exclusive 2^nd^ restriction site. These gene cassettes were cloned into a vector, colony-amplified and singularized. All SIRV sequence in the purified plasmids were verified by Sanger-sequencing to identify the correct clones. The E. coli cultures were grown in batches to obtain plasmids in the lower µg-scale. Double digestion of isolated plasmids with XhoI and NsiI showed correct insert size and complete restriction. Linearized, silica-purified plasmids served as templates in in vitro transcription reactions using T7 transcription kits (AmpliScribe T7 High Yield Transcription Kit, and AmpliScribe T7 Flash Transcription Kit, Epicentre, Madison, WI). The DNase-treated, phenol-extracted and silica-purified in vitro transcription products were assessed for concentration and purity by spectrophotometry (NanoDrop, Thermo Fisher Scientific, Waltham, MA) and for integrity by capillary electrophoresis (2100 Bioanalyzer, RNA 6000 Pico Kit, Agilent Technologies, Santa Clara, CA).

In the context of variant verification RNA integrity is a very important measure. Fragments arising from incomplete transcription might impose errors on the correct determination of variants which share those sequences and thereby also affect the overall gene coverage. The integrity of the transcription products was very heterogeneous as expected, given the broad sequence variation and the length of the SIRV transcripts (average 1.1 kb with 14 RNAs between 2.0 and 2.5 kb). A majority of transcripts therefore had to be purified by at least one of two purification methods. Purification method PP1 is selective for poly(A)-tails and neglects prematurely terminated transcription products. Purification method PP2 is based on size-selective quantitative electrophoresis separating the correctly sized main products from shorter fragments (transcription break off and degraded products) and longer fragments (run-through transcription products). After purification, the 69 SIRV RNAs were assessed to be on average 7.36±3.43 % in the pre-peak fraction, 90.31±3.72 % in the main-peak fraction (corresponding to RNAs of correct length), and 2.36±3.04 % in the post-peak fraction (mass percentages, wt%). Noteworthy, the risks of a beginning degradation during the initial denaturation (2 min at 70°C) and the not fully denaturing conditions during RNA microcapillary electrophoresis, tiny noise levels in the baseline, and slightly different shapes of the main peak result in a rather over-than underestimation of pre-and post-peak fractions.

rRNasin (Promega, Madison, WI) was added to 0.8 U/µl to minimize the risks of RNase contamination induced RNA degradation. The SIRVs were quantified by absorbance spectroscopy to adjust all stock solutions to a base concentration of close to, but above 50 ng/µl. RNA purity was very high with A _260/280_ 2.14±0.12, and A _260/230_ 2.17±0.20.

### SIRV Mixing

SIRV mixes were designed that contain pools of SIRVs (SubMixes) in defined amounts and ratios by combining accurate volumes of the stock solutions in sufficiently large batch sizes. Pipetting errors vary depending on transfer volumes, and range from ±4 % for 2 µl to ±0.8 % for >100 µl transfers. The precision was experimentally determined with blank solutions. Starting with the stock concentration measurement (NanoDrop accuracy ±2 ng/µl for 50 ng/µl, or ±4 %), and accounting for the entire mixing pathway, the accumulative concentration error is expected to range between ±8 % and ±4.7 %. Therefore, in the data evaluation one has to account for the experimental fuzziness by allowing for lower accuracy thresholds of ±8 % on the linear scale, or ±0.11 on the log_2_-fold scale, respectively. The SIRV concentration ratios between two mixes are more precise because only one final pipetting step defines the concentration differences and synchronizes all SIRVs, which belong to the same SubMix. Here, a maximal error of ±4 % (or ±0.057 on the log_2_-fold scale) can be expected between SubMixes, while all SIRVs of the same SubMix must propagate coherently into the final mixes. Bioanalyzer traces were used to monitor the relative propagation of the SIRVs, PreMixes and SubMixes during the mixing (Fig. 2B). In addition, the accurate pipetting of the 8 PreMixes was controlled by checksums of Nanodrop concentration measurements which deviated by only 0.002±3.4 % from the calculated target concentrations, and by weighing on an analytical balance, which showed a deviation of 1.8±0.65 %. The final concentration of the SIRV mixes are set to be 25.3 ng/µl.

### Spiking of Reference RNA with SIRV Mixes

In general, SIRV mixes can be used with crude cell extracts, purified total RNA, rRNA-depleted RNA or poly(A)-enriched RNA. Important, the spike-in ratios have to be chosen in concordance with the desired final SIRV content, the RNA preparation method and, if known, the expected mRNA content. One widely applicable spike-in ratio can be obtained by adding 0.06 ng of controls to approximately 100 ng of total RNA. In the presented examples, the total RNA mass of one single aliquot RNA was 100 ng of UHRR in RC-0, and 112.8 ng of HBRR in RC-1. The reference RNA in RC-2 was calculated to be 104.27 ng (Fig. S2). The high-concentrated and thereby stable SIRV mix stocks with 25.3 ng/µl, and ERCC mix stocks with 30 ng/µl (ERCC Spike-In Mixes 1 and 2 by Ambion, Thermo Fisher, Waltham, MA) were diluted just prior to the spiking to facilitate the mixing of manageable volumes. For example, the final 1:1000 dilution reduced the SIRV mix concentration to 25.3 pg/µl from which 2.37 µl were required to spike a 100 ng total RNA sample with 0.06 ng SIRVs. The RC-samples were produced in large batches to yield hundreds of samples allowing for a protocol with fewer dilutions and larger volumes reducing thereby sources of experimental errors.

### NGS Library Preparation and Sequencing

The experiments were carried out in technical triplicates. From 1 µg RC-0, RC-1 or RC-2 RNA in total nine NGS libraries were prepared by Fasteris SA (Plan-les-Ouates, CH) using the TruSeq stranded mRNA library preparation kit with polyA selection (Illumina Inc., San Diego, CA). All primary library templates were amplified with 14 PCR cycles to reach reproducible concentrations, 155±45 nM, and insert sizes, 165±10 bp. An equimolar lane-mix was sequenced at the CSF Campus Science Support Facilities GmbH (CSF, Vienna, AUT) in PE125 mode in one lane of a HiSeq 2500 flowcell in 125 bp paired-end mode using V4 chemistry. The run yielded 305 M raw reads, 293 M after demultiplexing and trimming which corresponds to a read depth of 32.6±2.2 M reads per library.

### Read Processing and Mapping

Standard data processing routines (bbduk, part of BBMap by A Bushnell to be found at: sourceforge.net/projects/bbmap/) were applied removing low quality reads and trimming adapter sequences. Due to variable stringencies the mappers produced different read statistics: STAR mapped against the genome 89.7±5.9 % of the reads (86.4±5.6 % uniquely), and TopHat2 85.2±5.7 % (79.4±5.2 % uniquely). Bowtie2 mapped against the transcriptome 69.3±1.8 % reads (15.1±1.5 % uniquely).

### Estimation of the mRNA Contents

Based on the obtained read distributions the relative mass partition between external controls and endogenous RNA fraction can be estimated. Because the library preparation uses a poly(A)-enrichment step the targeted endogenous RNA corresponds almost exclusively to the mRNA content of the samples. The propagation of the input mass ratio to the output read ratio depends on both the mRNA content and the efficiencies of how controls and mRNA proceed through the library preparation. The relationship is described by Eqn. 1.

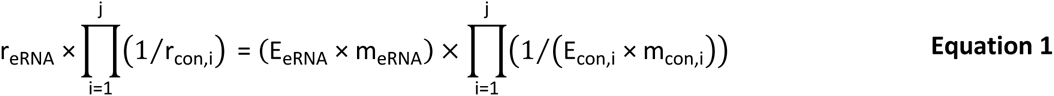

with r, number of reads; from eRNA, the endogenous RNA fraction, here mRNA, and con, the controls, here SIRVs and ERCC; E, the relative efficiency of how the fraction proceeds through the workflow; and m, the input mass.

Ratios are measured as relative read counts, and the results as a product of efficiencies and apparent amounts are also relative measures. Therefore, at least one de facto arbitrary reference point needs to be defined to deduce comparable, absolute results. Several reference points or default values can be defined which lead to different dependent variables, and imply slightly variable interpretation of the results. The following three calculation examples are simple deductions from the ratio shown in Eqn. 1 with the ERCC (con,1), and the SIRVs (con,2) respectively. Variation 1 is explained in greater detail, while in the other variants only the differing aspects are highlighted.

#### Variation 1

i. Here, the first assumption is that the mRNA content in UHRR (Agilent, Santa Clara, CA) is 3 % (factor 0.03), and 2 % (factor 0.02) in HBRR (Ambion, Thermo Fisher, Waltham, MA), which stays in close agreement to previous estimates concluded from experiments by Shippy et al. in 2006. The amounts of total RNA were measured by Nanodrop absorbance spectroscopy. The UHRR samples contained 100 ng and the HBRR samples 112.8 ng. The mixing ratio of (2× RC-0): (1× RC-1) determines the concentration of the 3^rd^ sample, RC-2, and further the amount of total RNA (104.27 ng) as well as its related mRNA content, (2×100 ng × 3 % + 1×112.8 ng × 2 %) / (2×100 ng + 1×112.8 ng) = 2.64 % (factor 0.026). The mRNA masses of the samples are calculated by the measured total RNA input amounts and default mRNA contents are 3 ng, 2.256 ng, and 2.752 ng. The second assumption is that the endogenous mRNAs proceed with the maximal efficiency of 100 % (factor 1) through library preparation and sequencing. The input of the spike-in controls is the product of the concentration values which were either measured as in the case of the SIRV stocks, or stated by the vendor as in the case of the ERCC stocks, and the pipetted volumes. Both are treated as de facto measured values.
ii. The relative amounts of reads which have been assigned to either endogenous mRNA, ERCCs, or SIRVs are obtained from the mapping statistics (read classification).
iii. The ratio of mass input to reads output allows to calculate the propagation efficiency of the controls relative to the default efficiency of the endogenous mRNA. In the present experiment, the SIRVs progress with a relative efficiency of 86.7±5.7 % and the ERCCs with 41.6±4.7 %. The significant difference between the relative efficiency of both controls is very likely caused by the different lengths of the poly(A)-tails which might influence a different binding efficiency during poly(A)-enrichment step. The SIRVs with a 30 nt long poly(A)-tail perform much more in concordance with the endogenous mRNA as the ERCCs which contain shorter and variable poly(A)-tails (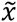, 23 nt). The observed small differences in the experiments using RC-0, RC-1, or RC-2 could be caused by a two-way interference of samples and controls. However, the deviations are small and can rather be a result of minor inaccuracies in determining the relative input amounts. Both hypotheses would need to be verified by a series of repeats with the aim to resolve potential influences of pipetting during the sample preparation, variations in the library preparation, and other contributions.

#### Variation 2

Here, no varying interferences between samples and controls are permitted which means that relative efficiencies of the controls are considered to be constant. The causes of the experimental variations are exclusively assigned to the input amounts (alike hypothesis two in the above example in discussing the deviation of the results in RC-2). This would translate into input standard deviations of 11.8 % (SIRVs) and 7.6 % (ERCCs). The variations meld all accumulating errors made in determining the concentration of the control stock solutions, preparing the subsequent dilutions, pipetting the samples and controls during the spike-in step itself, as well as determining the amount of sample RNA in the first place. Against the background of potential errors the calculated results can be considered as highly consistent.

#### Variation 3

In addition to the assumptions made for the measurement of sample RC-0 the spike-in of 0.06 ng SIRVs to all samples is also set as default. Alike variation 2 the experiments in RC-1, and RC-2, adopt the same default setting for the relative efficiencies calculated in RC-0.

According to the ratio of Eqn. 1 one can obtain mRNA contents as well as ERCC input amounts, in this case for samples RC-1, and RC-2. The mRNA content of the HBRR sample (RC-1) relative to the default mRNA content in the UHRR sample (set to 3 %) would be 1.72 %, lower than the previously assumed 2 %. According to the present mixing scheme it would also imply that sample RC-2 has an mRNA content of 2.54 %, up from the calculated 2.34 %. The offsets of −7.9 % is again small and very likely refers to the level of accuracy in the experimental results as discussed above. Using this method the spiking of samples with controls such as the SIRVs allows to closely estimate the relative mRNA content of any unknown sample, and to draw conclusions e.g. on the transcriptional state of tissues, cell suspension or individual cells.

### Coefficient of Deviation

We implemented a strand specific Coefficient of Deviation (CoD) which aims to reflect the quality of NGS pipelines targeted at specific effects, in particular, ‘spikeness’ of the coverage, while systematic effects like drop of the coverage at Transcription Start Sites (TSS), and Transcription End Sites (TES) are being suppressed. The CoD is implemented by Eqn. 2,

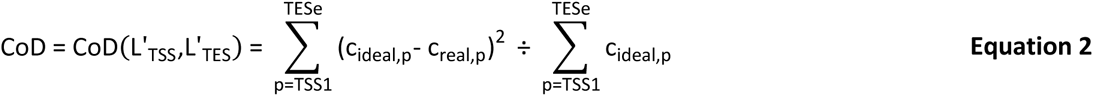

which is calculated in strand orientation for contigs along all exonic nucleotide position (p) from the first annotated TSS (TSS1) up to the last annotated TES (TESe) whereby c_ideal_ are the ideal coverages with the optimized transient regions (L’_TSS_, L’_TES_), and c_real_ are the real coverages scaled as such that the total integrals along all positions of ideal and real coverages are equal.

The transient function at TSS and TES is the error function, erf(x), which corresponds to Cufflinks assumption about normally distributed fragment lengths (Trapnell et al. 2012). Since erf(x) is a non-algebraic function we used the algebraic approximation by Abramowitz and Stegun (https://en.wikipedia.org/wiki/Error_function; eq. 7.1.25). Once the optima (L’_TSS_, L’_TES_) are found, we declare CoD(L’_TSS_, L’_TES_) to be the ultimate CoD value which mostly eliminated the transient region effects. We implemented an optimization algorithm (Nelder-Mead downhill simplex method) and used as starting values (*L^0^_TSS_*, *L^0^_TES_*) lengths of 150. In some cases, the optimum found can be beyond a reasonable range of values. Therefore, we censor the output L_opt_ of the mathematical optimization as follows, L_censored_ = 50 if L’ < 50; L_censored_ = 350 if L’ > 350. This corresponds to the maximal range of 200±150. This censoring needed to be applied in 8 cases out of 84 SIRV/mix/strand/end combinations, i.e. less than in 10 %.

### Transcript Concentration Measurements

In order to estimate expression values on transcript levels we applied RSEM (Li and Dewey 2011) which uses a Bayesian Network to compute the likelihood of each fragment belonging to a given transcript. Salmon (Patro et al. 2015) estimates the transcript expression values by a k-mer based mapping. Similar to RSEM, it uses an expectation-maximization (EM) algorithm to compute the maximum likelihood of relative abundances. Cufflinks2 (Trapnell et al. 2013) estimates the transcript abundances with a coverage-based maximum likelihood approach. It determines the probability of read assignments to isoforms.

### Differential Expression Analysis

We used the DESeq2 package (Love et al. 2014) for calculation of fold-differences between the results from different pipelines and different input samples mixtures. As explained in the main text, all outputs from the evaluated pipelines (in form of relative concentration measures) have been scaled. Scaling by linear transformation is needed as a consequence of the different mRNA contents in the samples but also to eventually compensate any potential pipetting errors when small aliquots, typically in the range of very few microliter, are added to individual samples. The RC-samples in this experiment were produced in a large batch to ensure high accuracy in volumetric ratios. The main sample pool is practically invariant to the small relative contribution of the controls but was technically treated in the same way by scaling the main sample pools to identical sizes. First, SIRV concentrations were scaled to reach together 69 in E0, 68.5 in E1, and 70.8 in E2, which is the dimensionless value identical to the fmol/µl in the respective SIRV stock solutions. Then, the lower threshold was set to 10^−6^, which corresponds to original FPKM values of around 5∙10^−3^, 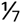 ^th^ of one read for a 1 kb long transcript at a read depth of 30 Mio reads, or 1 gene specific read to be divided between 7 equally possible transcript variants at said 1 kb length section. The threshold is to be understood as less than, or equal to, 10^−6^ which defines an averaged read resolution of the particular workflow at the given read depth. By these means all ratios remain defined. Second, for the DESeq2 calculations we multiplied all values with 10^6^, after which fractions were rounded to the closest integer. We understand that for no-change hypothesis testing, which is the main purpose of the DESeq2 package, transforming the relative concentrations (FPKM or TPM) to ‘quasi-read-counts’ is not recommended, since DESeq2 internally corrects for library size by using size factors. However, in our application, we do not use the probability of the hypothesis rejection (P values) but the raw fold changes detected (irrespectively of the dispersion values for that given transcript). We also took into account the potential danger of censoring the input (by unintentionally eliminating transcripts represented by extremely low FPKM or TPM values) by setting a suitable threshold as described above. The figures show only two differential expression analyses because the 3rd concentration ratio, E2/E0, is the product of E1/E0 and E2/E1.

### Error Rate Analysis

The four red data points, two from the SIRV controls (circles) and two from the endogenous mRNAs (squares), from the best agreement (P3 vs, P1), and the four corresponding black data points from the worst agreement (P4 vs. P2) as shown in fig. 5E were fitted by linear approximation using the relationship as derived for Eqn. 3,

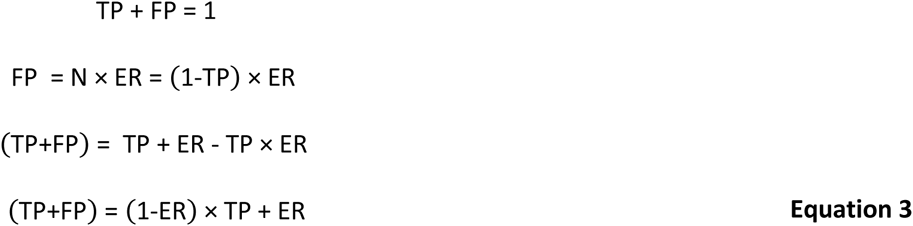

where TP, is the relative portion of true positives (differentially expressed as in our definition possessing a LFC > 1); N, the relative portion of negatives; ER, the error rate at which negatives are falsely detected as positives to result in false positives, FP. The least squares fitting method was applied to derive values for the slope (1-ER), and the intercept ER.

The transformation of Eqn. 3 leads to the relationship of Eqn. 4,

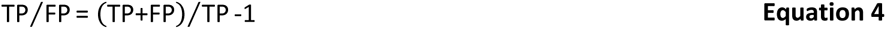

which has been used for the graph in fig. 5F. In the present context, the concept of ROC-curves is not applicable because the true or false negatives are indistinguishable as long the ground truth of the endogenous RNA are unknown.

## Data Access

Sequencces and source data files data are in the process to be deposited at NCBI under BioProject ID: PRJNA312485.

## Acknowledgements

We thank S.A. Munro and M. Salit (Material Measurement Laboratory, National Institute of Standards and Technology) for their efforts in further developing the concepts of the External RNA Control Consortium (ERCC) in an updated suite of RNA controls and analysis methods called ERCC 2.0. We thank A. Tuerk for his contributions in finding suitable human model genes, G. Wiktorin for carrying out the RSEM calculations, M. Mihaylova in helping with the graphics design, and I. Gabler for critically reading the manuscript.

## Author Contributions

L.P., P.K. and T.R. designed the sequences and mixtures. P.K. and G.H. prepared the in vitro transcripts and spike-in mixtures. T.R., A.S. and L.P. designed the research and the experiments. M.A., I.H. and T.R. evaluated the data. T.R. and L.P. wrote the manuscript.

